# Reducing ploughing promotes ground-nesting flying insects

**DOI:** 10.1101/2025.10.09.680943

**Authors:** Christopher Hellerich, Michael Garratt, Alexandra-Maria Klein, Felix Fornoff, Anne-Christine Mupepele

## Abstract

Many flying insects, such as bees, wasps, and hoverflies, live aboveground, but depend on soil for nesting, development, and overwintering. In agricultural landscapes, soil is managed for production and therefore frequently subjected to disturbances such as ploughing. The impacts of ploughing on flying insects that spend part of their life cycle belowground remain largely unknown.

To investigate the effects of ploughing on ground-nesting flying insects, we conducted a two-year field experiment in flower strips subjected to different treatments, ranging from annual ploughing to four years without ploughing. Insects emerging from the soil were sampled using emergence traps, allowing a direct assessment of their response to ploughing at different frequencies. For each treatment, we measured insect biomass, abundance, and body size.

We found that ploughing substantially reduced flying insect biomass. When sites were left unploughed, biomass increased rapidly, particularly during the first years of recovery. However, regardless of the time since the last disturbance, ploughing always reduced insect biomass to similarly low levels, driven primarily by declines in the abundance of large insects. Our findings highlight that even moderate reductions in ploughing frequency, for example, only every second year, can benefit ground-nesting flying insects and point to the potential for incorporating reduced ploughing frequencies into agricultural management and agri-environmental schemes.

## 1. Introduction

Insects are integral to the productivity of global agricultural systems: By providing a multitude of ecosystem services - such as pollination, nutrient cycling or pest control - they contribute directly to agro-economical output and are essential for safeguarding global food security (Dainese et al., 2019; Klein et al., 2007; Losey & Vaughan, 2006; Requier et al., 2022). Conserving insect diversity and abundance is therefore not just an ecological priority, but a requirement for sustainable agricultural production. Meeting this requirement depends on managing agricultural landscapes with viable and diverse habitats that provide foraging, nesting, and overwintering resources for insects (Gebert et al., 2024; Pimentel et al., 1992).

Agricultural land is often shaped by management practices, such as fertilizing or ploughing. The intensification of these practices is considered to be a major driver for biodiversity loss (Sponagel et al., 2025; Tscharntke et al., 2005). Ploughing has a strong impact on soil, which is turned over to prepare the seedbed for new cultures or to manage weeds (Soane et al., 2012). Conventional ploughing is usually carried out annually, or sometimes more than once per year. When the ploughing frequency is reduced, it is generally referred to under the broader concept of ‘conservation tillage’ (Liang et al., 2025), which ranges from biennial to minimum or even zero tillage. In any case, each ploughing event interrupts the natural succession of plants and alters the structure and layering of the soil (Rowen et al., 2020). Meanwhile, soil plays an important role in the life cycle of many insects, serving as a site for reproduction (i.e., nesting), overwintering (i.e., diapause), and larval development (Lavelle & Spain, 2005). Ploughing disturbs soil and can have detrimental effects on ground-dwelling insects, with variation depending on taxon and body size (Ganser et al., 2019; Kladivko, 2001; Pfiffner & Luka, 2000).

Current knowledge about the effects of disturbance on soil fauna is mostly limited to specific taxa due to the employed sampling method (Moreno-García et al., 2024; Rowen et al., 2020). Soil cores or pitfall traps predominantly target epigean and soil-dwelling taxa of the meso- and macrofauna, such as springtails, nematodes, earthworms or carabid beetles (Müller et al., 2022; Wardle, 1995; but see Pfiffner & Luka, 2000). Most of these species cannot fly and are limited in their capacity to actively disperse and colonize new areas. Using other sampling methods such as sweep netting or pan trapping to sample flying taxa (e.g., Hymenoptera, Coleoptera or Orthoptera) can only detect indirect effects of soil disturbance, for example, those mediated through changes in soil litter structure or weed density and diversity (Stinner & House, 1990). They do not allow a direct assessment of emergence from soil and whether sampled insects are using a particular habitat for nesting (Revilla-Martin et al., 2025; but see Ullmann et al., 2016).

In the present study, we used emergence traps (‘e-traps’) in a highly replicated field experiment to investigate the direct effects of spring ploughing and its frequencies on the biomass, abundance, and body size of ground-nesting flying insects in perennial flower strips. Insect biomass can be a suitable proxy for the state of an insect community and its capacity to provide ecosystem services, because it reflects abundance, and to a limited extent also species richness (Hallmann et al., 2021; Winfree et al., 2025). The taxa we targeted spend only parts of their life cycle below ground but disperse, feed, and mate aboveground, typically by means of flying. Because of their increased mobility, they play a key role in ensuring the provision of aboveground ecosystem services such as pollination and pest control (Kremen et al., 2007), the quick recolonization of disturbed habitats (Lundberg & Moberg, 2003) and, as a consequence, the ecological resilience in agricultural landscapes (Tscharntke et al., 2005).

The goal of our research was to assess the impact of a) ploughing, and b) reduced ploughing frequency, on ground-nesting and ground-overwintering flying insects, in terms of biomass, abundance, and body size.

Based on previous findings that ploughing negatively affects many soil-dwelling arthropod taxa (Müller et al., 2022; Wardle, 1995), we hypothesized that (a) ploughing reduces the biomass and abundance of ground-nesting and ground-overwintering flying insects, with disproportionate effects on larger-bodied species than compared to smaller species (Andrén & Lagerlöf, 1983; Kladivko, 2001). (b) We further hypothesized that insect biomass increases when reducing ploughing frequency, i.e. time since last ploughing (Boetzl et al., 2022). We assumed that insect biomass increases mostly in the first year, with a decreasing rate thereafter, reflecting saturating community recovery (Schwerk & Szyszko, 2011) through the continuous provision of undisturbed nesting and overwintering habitat.

## 2. Materials and methods

### 2.1 Insect sampling

The study was conducted in 2023 and 2024 in the Upper Rhine Valley, Baden-Württemberg, Germany. The study region (∼200□m altitude) includes the flat Rhine plain with deep gravel soils and the hilly Tuniberg and Kaiserstuhl, shaped by loess deposits on volcanic bedrock. It has a sub-Atlantic climate influenced by maritime and continental air masses. With a mean annual temperature of ∼10□°C, it is among Central Europe’s warmest regions; monthly maxima often exceed 30□°C in summer. Annual precipitation ranges between 600–700□mm, with up to 70 frost days. Land use is dominated by intensive arable farming (e.g., maize, sugar beet, cereals, rapeseed) and various specialty or permanent crops (e.g., asparagus, strawberries, tobacco, vineyards, stone fruits, hops); grasslands are of minor importance.

We established our experimental sites in 2023 across 12 flower strips on three conventional farms. Flower strips covered an area of 0.13 to 0.55 ha (mean = 0.25, SD = 0.12) and were 0.03 to 17.23 km (mean = 8.77, SD = 7.09) apart from each other (see Suppl. 1 for site locations). The flower mixtures used are generally attractive to many insect groups, providing food resources (Boetzl et al., 2022; von Königslöw et al., 2022; Williams et al., 2024). Seven sites had already been managed under a flower strip scheme for up to three years, while the remaining five were still in arable production (corn or wheat). At the start of the experiment, new flower strip sites were ploughed using a conventional mouldboard plough, and then sown with a perennial wildflower seed mix, reflecting the flower strip establishment of the older flower strip sites. Each site was divided into an unploughed control plot and a ploughed treatment plot, but one newly established site was not ploughed due to miscommunication with the farmer, and thus only served as a control plot. The control plots were left unploughed and, in some cases, these were split in the second year into new ploughed and control plots. Ploughing of treatment plots was conducted between January and February 2024. Exceptional ploughing in May 2023 took place at two of the plots that had been established three years earlier, as part of the regular farm operations on one of the farms. This way, flower strips were ploughed after one, two or three years of establishment, depending on the year of establishment (see Suppl. 2 for a table of replicates by years of establishment and ploughing). In combination with control plots that had remained unploughed for one to four years, this research design enabled us to study the impact of a) ploughing, and b) reduced ploughing frequency, based on data collected over two consecutive sampling years.

We used custom-built emergence traps (e-traps) to sample ground-nesting insects on control and treatment plots. E-traps are a suitable passive method for non-specific sampling of insects at their place of nesting or overwintering. Here, we use the term ‘ground-nesting’ synonymous to ‘overwintering’ but acknowledge that collected individuals may have emerged from an overwintering place that is not necessarily a nest *sensu stricto* (i.e., the place of reproduction, see e.g. Westrich, 2018). Typically, the method is biased towards flying taxa due to the elevated position of the collection unit (Hellerich et al., 2025). While e-traps have been used for a variety of research questions related to arthropod nesting and overwintering (Boetzl et al., 2022), e-trap studies on ploughing effects are rare and often taxon-specific (Funderburk et al., 1983; Ganser et al., 2019; Hanson et al., 2015; Holland & Reynolds, 2003; Thorbek & Bilde, 2004).

E-traps consisted of conventional hiking tents from which the ground floor was removed and which covered ca. 2.2 m^2^ of ground per trap. Collection units were installed on top of the traps, each filled with 100 ml of a preservative solution (40% Ethanol, 10% Glycerin, odour-free dish soap) to kill and preserve specimens while minimizing evaporation loss (see Hellerich et al. (2025) for a full description of the e-trap method).

Eight traps were placed along two semi-random transects per plot. To ensure spatial representation, distances between traps depended on the plot length (Hellerich et al., 2025). In 2023, 160 e-traps were installed between April and May and remained at the same place for an average of 99 days (SD(n) = 19 days). In 2024, 208 e-traps were installed in March and were in place for an average of 128 days (SD(n) = 1 day). The number of traps varied between the two years, because two sites and four control plots were only sampled in 2024, while the two treatment plots that were ploughed in 2023 were discontinued. E-traps were emptied every second week (14.2 days, SD(n) = 1.5 days) by replacing the collection unit liquid.

### 2.2 Insect biomass, abundance, and body size data

Biomass of sampled insects was measured in a moist state to preserve samples for future analyses, following a standardized protocol (Hallmann et al., 2017). First, sampled insects were emptied into a stainless-steel sieve (1mm mesh size). The contents of the sieve were shaken out onto a paper towel and spread out to remove any adhering liquid. Any non-arthropod species (e.g., Gastropods) were removed from the sample. Subsequently, the still-moist sample was weighed to the nearest 0.001 g using an analytical precision scale (Kern ADJ 200-4) and subsequently stored in Scheerpelz solution (70% Ethanol, 5% acetic acid). To standardize for sampling effort, insect biomass data were divided by the number of days in each sampling round.

We observed episodic mass-occurrence samples dominated by a single species, typically clustered ant nuptial flights in July. Because these events represent short-term dispersal swarms rather than local ground insect dynamics, we treated them as a distinct ‘outbreak’ process. Outbreaks were defined as >100 individuals <5 mm or >50 individuals ≥5 mm of a single species; such cases were visually obvious outliers, with abundances orders of magnitude above typical samples. In total, 144 samples met this criterion and were excluded prior to modelling (retained n = 2,847). For transparency, we provide the models including all samples in the supplementary material (Suppl. 3b,c); treatment effects remained qualitatively similar, but variance-related tests became significant due to the influence of a few extreme values.

To analyse the effects of ploughing on insect abundance and body size, individuals of a subset of samples in 2024 were categorized by body size (small <5 mm or large ≥5 mm) using millimetre paper and counted separately. Body size was measured without antennae and ovipositors. Per sampling round, we measured and counted all individuals of two out of the eight e-trap samples of each plot (n = 336). Ten of these samples were excluded prior to the analysis due to mass-occurrences (see above). To standardize for sampling effort and to align with the insect biomass data, insect abundance data were divided by the number of days in each sampling round. Due to the expected high numbers of individuals, no systematic taxonomic assessments were performed, but the most abundant taxonomic groups were noted qualitatively to provide an overview of the sampled arthropods. These included taxa from various insect families, with only minor contributions from non-insect taxa like Araneae and Acari. Among the larger insects, ground-nesting or ground-overwintering Hymenoptera (e.g., Vespoidea, Apoidea, Ichneumonidae), Diptera (e.g., Limoniidae, Tipulidae, and various Brachycera) and Lepidoptera as well as several families of Coleoptera (e.g., Cantharidae, Curculionidae, Carabidae, Oedemeridae), Orthoptera, and Dermaptera were observed. For the smaller species, various above- and belowground nesting families of Diptera (e.g., Sciaridae, Drosophilidae), Hemiptera (mainly Sternorrhyncha such as Aphididae, and Auchenorrhyncha), and small Coleoptera and Hymenoptera were observed.

### 2.3 Statistical analysis

All analyses were performed using R 4.4.1 (R Core Team, 2024). The effects of ploughing and of reduced ploughing frequency on ground-nesting insect biomass were assessed using a generalized linear mixed model (hereafter called ‘insect biomass model’) in the ‘glmmTMB’ package (Brooks et al., 2017). To assess how ploughing affected insect biomass indirectly via changes in insect abundance and body size, a structural equation model (SEM) was fitted in the ‘piecewiseSEM’ package (Lefcheck, 2016) on a subset of data where individuals were counted.

#### Insect biomass model

The insect biomass model included daily insect biomass as the response variable (see Suppl. 3a for the model term and diagnostics). Predictors (fixed effects) were the time since establishment (in years), ploughing, sampling year, and Julian day. ‘Trap’ nested within ‘sampling site’ were included as nested random factors. The insect biomass model was fitted using a Gamma-distribution with a log-link. The best model was selected using residual diagnostics of the ‘DHARMa’ package (Hartig, 2024), Akaikes information criterion (AIC) and likelihood-ratio-tests of the ‘stats’ package (R Core Team, 2024). Model performance was assessed using the root-mean square error (lower RMSE indicated better accuracy) and marginal and conditional R^2^ in the ‘performance’ package (Lüdecke et al., 2021). Based on our hypothesis of a saturating recovery with time since establishment, we modelled this predictor using an inverse function (1 / time since establishment). While inclusion of the inverse function did not significantly improve model performance (Δ RMSE < 0.001), residual diagnostics indicated better model fit: the model with the linear function showed significant deviation from residual uniformity (DHARMa KS test), whereas the model with the inverse function did not. Similarly, because seasonal dynamics often exhibit unimodal patterns (Forrest & Miller-Rushing, 2010), we modelled Julian day as a quadratic polynomial term, which significantly improved model fit (‘Chisq’-test: *p*=<0.001) compared to linear alternatives.

#### SEM

The SEM model followed a bottom-up pathway with ‘ploughing’, ‘Julian day’ and ‘time since establishment’ influencing the top response ‘daily insect biomass’ directly and indirectly through their effects on the two mediating variables ‘daily abundance of small insects’ and ‘daily abundance of large insects’. The SEM comprised three component models: two separate models for each mediating variable and a third model for the top response variable (see Suppl. 3c for R model terms). To account for both direct and indirect effects, the top model included all upstream predictors and mediators as explanatory variables, except for ‘time since establishment’, which was dropped from the main model to avoid overparameterization (i.e., a saturated model with zero degrees of freedom). Its effects were captured through the mediator models. All component models included ‘trap’ as a covariate. ‘Site’ was only included as covariate in the small insects abundance model, where it improved model performance (Δ AIC >3). The top response model (daily insect biomass) was fitted using a gaussian distribution on the log-transformed response; due to the presence of zeroes, the two response variables for the small and large insect abundance models were log1p-transformed to fit a gaussian distribution in each model. All numeric variables were centred around their mean at zero (‘scale’-function of the R ‘base’ package) to ensure interpretability of parameter estimates; the factorial ploughing variable (0 = unploughed, 1 = ploughed) was transformed to numeric values to enable inclusion in the SEM but left unscaled to preserve its categorical interpretation and maintain reference-level meaning. A covariance path between daily large and small insect abundance was added to the SEM to account for their non-independence.

#### Post-hoc analysis

For the post-hoc analysis following model fitting, we differentiated between ‘initial ploughing’ which occurred to establish the flower strips, and ‘within-experiment ploughing’, which we defined as ploughing events that were conducted after one to three years of flower strip establishment to assess the effect of ploughing. The year of initial ploughing served as the baseline to assess the effects of reduced ploughing frequency (time without ploughing since flower strip establishment). Separating within-experiment ploughing ensured that comparisons to control plots were restricted to sites that were flower strips prior to the ploughing treatment, thereby avoiding it confounding with other land-use types (e.g., arable use before flower strip establishment). Pairwise contrasts were computed using the ‘emmeans’ package (Lenth, 2017), allowing estimation of percentage changes between control and within-experiment ploughed plots and across different times since establishment.

## 3. Results

We analysed a total of 2847 e-trap biomass samples taken during the two study years. Daily insect biomass ranged between 0.00007 to 0.37 g (mean = 0.03, SD = 0.04). In 2024, we counted 63,507 individuals in the subset of samples (n = 326), of which 59,007 (92,9%) belonged to the small group and 4,500 (7,1%) to the large group. Daily insect counts ranged between 0.07 to 145 insects (mean = 13.7, SD = 19.3).

### a) Effects of ploughing

#### Insect biomass

Spring ploughing negatively affected biomass of ground-nesting, flying insects (insect biomass model: z=-7.69, *p*=<0.001; *r*^*2*^_*m*_ = 0.37). Daily insect biomass was on average -62.51% (±9.99%) lower on within-experiment ploughed plots compared to non-ploughed control plots across time since establishment and season (see Figure 1b and Suppl. 3a for the model coefficient table).

**Figure 1.**
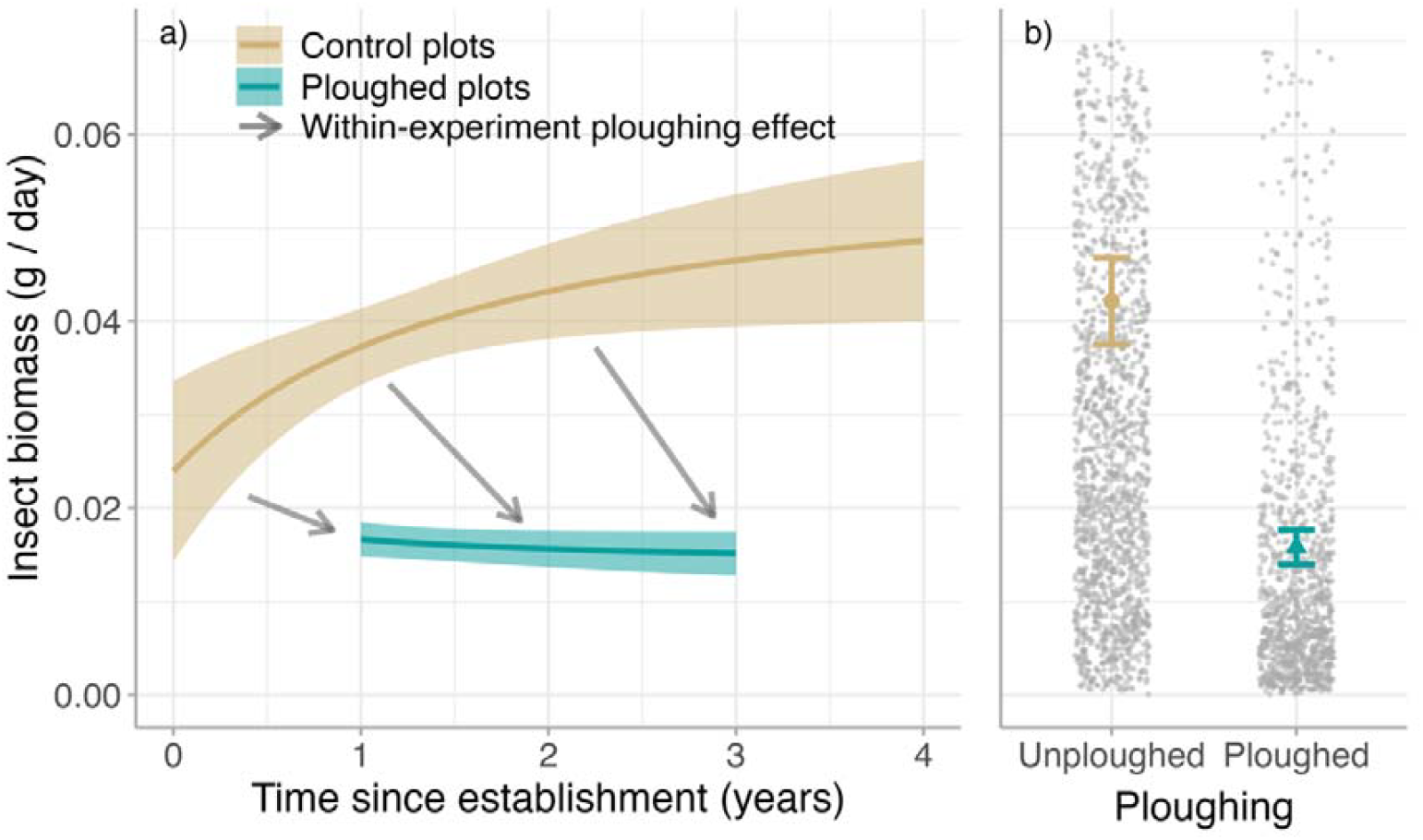
a) Insect biomass development after initial ploughing and effect of repeated within-experiment spring ploughing at different times after initial ploughing. Model predictions (lines) and standard errors of the mean (ribbons). b) Overall effect of within-experiment spring ploughing after 1, 2 and 3 years of flower strip establishment on insect biomass. Model predictions for unploughed (circle) and ploughed (triangle) plots, standard errors of the means (error bars) and underlying raw data (grey points)

#### Abundance and body size

The abundance of small and large insects equally contributed to the insect biomass (SEM: small insects: std. estimate = 0.456, ±0.032 SE, *p* =<0.001; large insects: std. estimate = 0.465, ±0.032 SE, *p* =<0.001; *R*^*2*^_*m*_ = 0.78; see Figure 2 and Suppl. 3c for the model coefficient table). The observed reduction of daily insect biomass after ploughing was only mediated by the significant reduction in the daily number of large insects (SEM std. estimate = -0.439 ±0.123 SE, *p* =<0.001; *R*^*2*^_*m*_ = 0.32; see Figure 2). The model didn’t indicate an effect of ploughing on small individuals (std. estimate = -0.149, ±0.16 SE, *p*=0.06; *R*^*2*^_*m*_ = 0.14), but ploughing had a direct negative effect on daily insect biomass (std. estimate = -0.094, ±0.085 SE, *p*=0.026, *R*^*2*^_*m*_ = 0.78).

**Figure 2.**
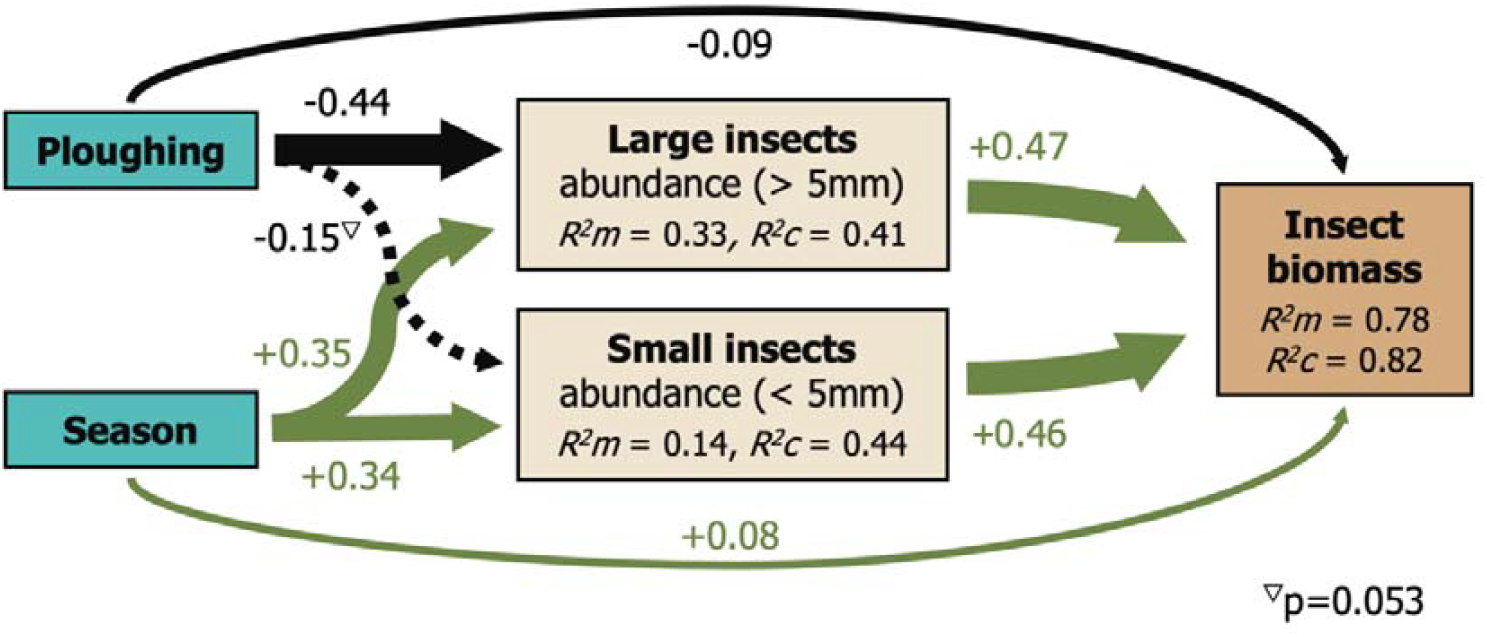
Structural equation model (SEM) showing positive (green) direct and indirect effects of season via large and small insect abundance on insect biomass and direct and indirect negative (black) effects of within-experiment spring ploughing on insect biomass. Ploughing reduced large insect abundance which indirectly reduced insect biomass. The amount of variation explained by each model is indicated with the *R*^*2*^ values. The non-significant effect of ploughing on small insect abundance is shown as a dashed line. The width of the arrows corresponds to the effect strength

#### Seasonal effect

The effect of spring ploughing on the daily biomass of soil-emerging insects significantly increased over the season (insect biomass model: estimate (linear term) = 10.2, ±1.98 SE, z=5.16, *p*=<0.001). Insect biomass accumulated quicker on control plots than on ploughed plots (see Figure 3), highlighting the persisting, increasing effect of spring ploughing throughout the year. Time of the year positively influenced both daily large insects abundance (SEM: std. estimate = 0.353, ±0.043 SE, *p*=<0.001) and daily small insects abundance (SEM: std. estimate = 0.339, ±0.042 SE, *p*=<0.001; see Figure 2).

**Figure 3.**
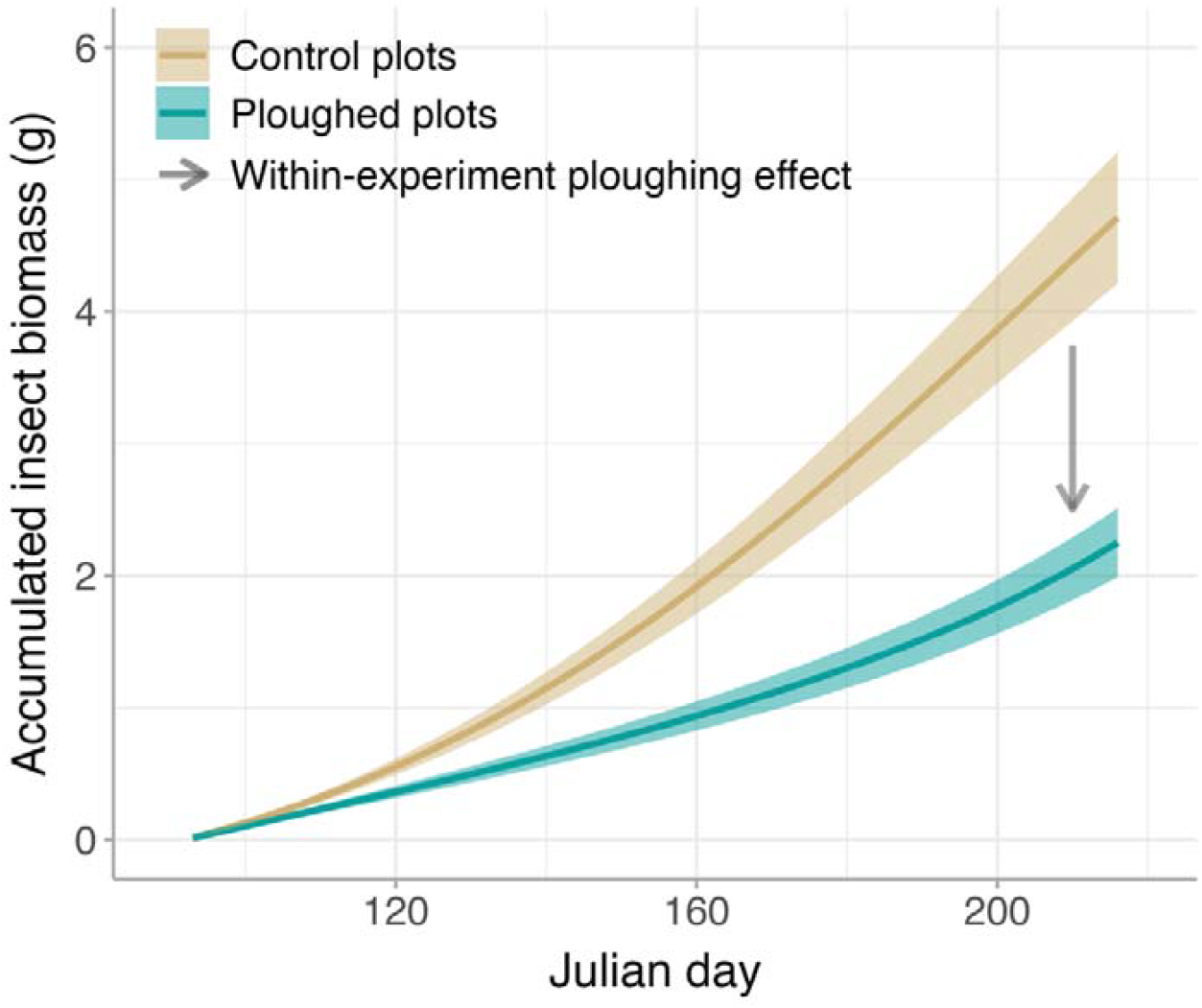
Accumulation of insect biomass over the season on control plots and on within-experiment spring ploughed plots. Model predictions (lines) and standard errors of the mean (ribbons)

### b) Effects of ploughing frequency

#### Insect biomass

Leaving sites unploughed over several years significantly increased daily insect biomass (insect biomass model: z=-3.75, *p*=<0.001). Daily insect biomass increased on unploughed plots progressively over time. However, the increase was strongest in the first years (year 1: +71.53% ±32.02%) and is expected to reach a plateau with no further increase after more than four years without ploughing (see Figure 1a). Due to the significant positive interaction with sampling year, the effect of time since establishment was largely offset in 2024 (z=3.51, *p*=<0.001). In this year, overall insect biomass was significantly lower than in 2023 (z=-6.58, *p*=<0.001).

#### Ploughing effect

The time since establishment significantly increased the ploughing effect (z=3.35, *p*=<0.001), which was higher on older sites (year 3: -67.42% ±8.22%) compared to younger sites (year 1: -55.33% ±7.24%, see Figure 1a). Spring ploughing always ‘reset’ insect biomass to the same low level compared to plots left unploughed, and the ploughed curve (Figure 1a) thus resembles the circumstances in intensive agricultural fields which are ploughed annually.

#### Abundance and body size

Time since establishment did not affect the daily abundance of either small or large insects in 2024 (SEM: small insects: *p*=0.706, *R*^*2*^_*m*_ = 0.14; large insects: *p*=<0.216, *R*^*2*^_*m*_ = 0.32; see Figure 2).

## 4. Discussion

Conventional mouldboard spring ploughing consistently reset ground-nesting flying insect emergence in flower strips, regardless of the time since the last spring ploughing. Larger insects (≥5 mm) were most strongly affected. A reduction in spring ploughing frequency increased insect emergence, particularly later in the season. To enhance insect abundance and support ecosystem service provision, ploughing should be minimised, and where ploughing is necessary, negative effects can be mitigated to a large extent by reducing frequency to be at most biennial (every two years).

### Effects of ploughing on insect biomass, abundance, and body size

Ploughing of flower strips effectively reset the biomass of ground-nesting flying insects at any time, even after several years without ploughing. Spring ploughed plots always exhibited equally low levels of insect biomass after ploughing, highlighting its profound negative direct effect on flying insects that were present in the soil in adult or preimaginal stage during ploughing. While previous studies have found evidence of direct negative effects of ploughing on organisms adapted to living entirely in or on the ground, such as earthworms or carabid beetles (Wardle, 1995), our findings demonstrate that insects with life cycles typically including a flight stage as well as a below ground stage, such as bees, flies, wasps, and moths, are affected in a comparable way, underscoring that ploughing can disrupt a wide range of insect functional groups beyond strictly soil-dwelling taxa. Our observed reduction in insect biomass following annual ploughing (-55%) also closely matches previous results by Ganser et al. (2019), who reported a similar reduction in overwintering epigeous and flying arthropod abundance after ploughing of annual wildflower strips (-59%). Because our SEM model indicated a strong link between insect abundance and biomass (see Figure 2), the close similarity between findings reinforces evidence for the negative impact of ploughing on ground-nesting insects and suggests that insect biomass is a suitable proxy for insect abundance (Gebert et al., 2024; Vereecken et al., 2021).

At initial flower strip establishment - which followed site ploughing to prepare the seedbed - insect biomass showed higher variance compared to plots that were ploughed after one, two or three years of establishment. This contradicts the idea that ploughing always ‘resets’ insect biomass to the same level but indicates that previous land use and management and the resulting species assemblages may have influenced the effect of ploughing (Hendrix et al., 1986; House & Alzugaray, 1989). A potential explanation was given by the SEM, which showed that both small and large insect abundance contributed equally to insect biomass on sites that had remained unploughed for more than one year. Annually ploughed sites however may host a larger abundance of smaller species at lower trophic levels that are adapted to the level of disturbance generated by ploughing (Andrén & Lagerlöf, 1983; House & Alzugaray, 1989; Kennedy et al., 2013). Life history traits such as short generation times or resistant resting stages may make them less vulnerable to - or even benefit from the disturbance (Andrén & Lagerlöf, 1983; Wardle, 1995). Our SEM also highlighted the importance of larger insects on older sites by showing that the negative effect of ploughing on insect biomass was mediated through reduced abundance of large insects, also supporting previous findings that larger species are more vulnerable to ploughing (Kladivko, 2001; Müller et al., 2022; Wardle, 1995). In addition, SEM results revealed a direct effect of ploughing on insect biomass independent of insect abundance (see Figure 2), indicating that ploughing also lowered the mean body weight of remaining individuals. Ploughed plots therefore may have contained proportionally more smaller insects, a size shift that was not captured by our broad size categories.

### Insect responses to reduced ploughing frequency

Many studies have examined the immediate effects of ploughing, or its effects several years later (Boetzl et al., 2022; Ganser et al., 2019; Wardle, 1995), yet little is known about the dynamics of soil fauna development in the years after ploughing. We observed the strongest increases in insect biomass particularly within the first few years of flower strip establishment. Our results indicate that even a moderate reduction in ploughing frequency by just 1-2 years can promote the rapid recovery of insect communities. Complementing previous observations of the complex effects of temporal continuity on ground-nesting, overwintering and succession of arthropods (Boetzl et al., 2022; Schwerk & Szyszko, 2011), our study especially shows that these effects may be non-linear and saturating, i.e. initially increasing after disturbance but then plateauing over time.

Besides the direct effect of ploughing through mechanical intervention, indirect effects may have contributed to the observed non-linear development of insect biomass. Ploughing can alter soil environmental conditions by relocating and reducing biomass, interrupting litter accumulation and access to food resources (Andersen, 2003; Andrén & Lagerlöf, 1983; Müller et al., 2022) or changing soil microclimate leading to drying out of the soil (House & Alzugaray, 1989). It can also cause alteration to soil food webs, for example by affecting slugs and earthworm abundance that serve as prey for predator species (Kennedy et al., 2013; Pelosi et al., 2009). Such consequences may be only present immediately after ploughing (e.g., bare soil), or diminish quickly within the following years (e.g., prey availability). In contrast, indirect effects may have also contributed to the rapid increase of insect biomass in the first year after ploughing by promoting influx and establishment of more mobile species which can colonize bare ground areas from adjacent sites rapidly (House & Stinner, 1983).

The increasing magnitude of the ploughing effect over time, along with the strong influence of large insects on the ploughing effect, suggests that older sites hosted comparatively larger, more plough-sensitive species. This pattern implies that both time since establishment and ploughing shape insect abundance as well as species composition (Kennedy et al., 2013). However, we did not find significant temporal changes in the abundance of either small or large insects. The strongest increase in insect biomass was observed during the first year after site establishment, but no new sites were established in 2024; thus, this effect was not represented in the data. Moreover, interannual variation in spring weather—particularly higher precipitation during peak emergence in April and May—likely contributed to annual variability in insect emergence. Together, the absence of first-year site effects and the influence of weather conditions, which may be strong drivers of insect biomass fluctuations (Müller et al., 2024), help explain the lack of temporal changes in insect abundance and the significantly lower overall biomass in 2024.

### Ecosystem service provision may benefit from reduced ploughing frequency across management types

The higher insect biomass observed on older sites, along with the strong negative effect of ploughing on biomass and abundance of large insects may be linked to changes in the provision of important aboveground ecosystem services, such as pollination, pest control or genetic dispersion (Kremen et al., 2007; Winfree et al., 2025). Most sampled taxa belonged to highly mobile groups such as Hymenoptera, Diptera, Lepidoptera or flying Coleoptera, which provide such services due to their dependency on aboveground food resources such as nectar and pollen of flowers, seeds, fruits or other insects (for predators and parasitoids). Various studies indeed suggest that reduced soil management can increase the numbers of natural enemies and pollinators (Appenfeller et al., 2022; Shuler et al., 2005; Stinner & House, 1990). Our findings thus indicate that annual ploughing not only suppresses insect biomass, but may also diminish the delivery of ecosystem services across agricultural landscapes.

We could show that spring ploughing has fundamental negative effects on ground-nesting flying insects, while reducing ploughing frequency can promote rapid, non-linear increases in insect emergence. The extent of these effects may vary between flower strips and arable fields, as insect occurrence is influenced by crop cover, agrochemical inputs, and management strategies (Appenfeller et al., 2022; Betancur-Corredor et al., 2022; Tschanz et al., 2023). Nevertheless, our findings suggest that the underlying direct and indirect mechanisms of ploughing remain consistent across management systems. As a result, similar ecological responses emerge even when site-specific conditions differ (Pfiffner & Luka, 2000). Besides underscoring the importance of maintaining undisturbed habitats within agricultural landscapes to support ground-nesting insects and associated ecosystem services, our findings highlight the ecological benefits of even short-term ploughing abatement (1–2 years), emphasizing the potential of short term, flexible and still effective measures in agri-environmental policy and management.

## Author contributions

**Christopher Hellerich**: Conceptualization, Methodology, Formal analysis, Validation, Investigation, Data Curation, Writing - Original Draft, Visualization, Project administration **Michael Garratt**: Conceptualization, Writing - Review & Editing, **Alexandra-Maria Klein**: Conceptualization, Resources, Writing - Review & Editing, Project administration, Funding acquisition **Felix Fornoff**: Conceptualization, Methodology, Validation, Writing - Review & Editing, Project administration **Anne-Christine Mupepele**: Conceptualization, Methodology, Validation, Writing - Review & Editing, Project administration, Funding acquisition

## Declaration of generative AI and AI-assisted technologies in the writing process

During the preparation of this work the authors used ChatGBT (OpenAI) in order to improve the language, grammar, spelling and overall readability. After using this tool, the authors reviewed and edited the content as needed and take full responsibility for the content of the published article.

## Acknowledgements

We gratefully acknowledge the financial support of the Ministry for the Environment, Climate and Energy Sector of Baden-Württemberg. We extend special thanks to the lower nature conservation authorities in Freiburg and Emmendingen for granting the necessary permits, and to the Regierungspräsidium Freiburg (Ref. 56 and 33), NABU Freiburg and all involved farmers for their support in the selection and management of the study sites.

## Data availability

Full model equations and parameters can be found in the supporting information. Data and R-code available private for review on figshare: https://figshare.com/s/ca3c29543b6e0f15642c

