## Supporting Information for "Reducing ploughing promotes ground-nesting flying insects"

*shared last authorship

This document includes:

### Suppl. 1: Map of the study sites

**Figure S1: The 12 study sites (blue triangles) were located in the south-west of Germany, in the Upper Rhine valley west of Freiburg im Breisgau (orange circle).**
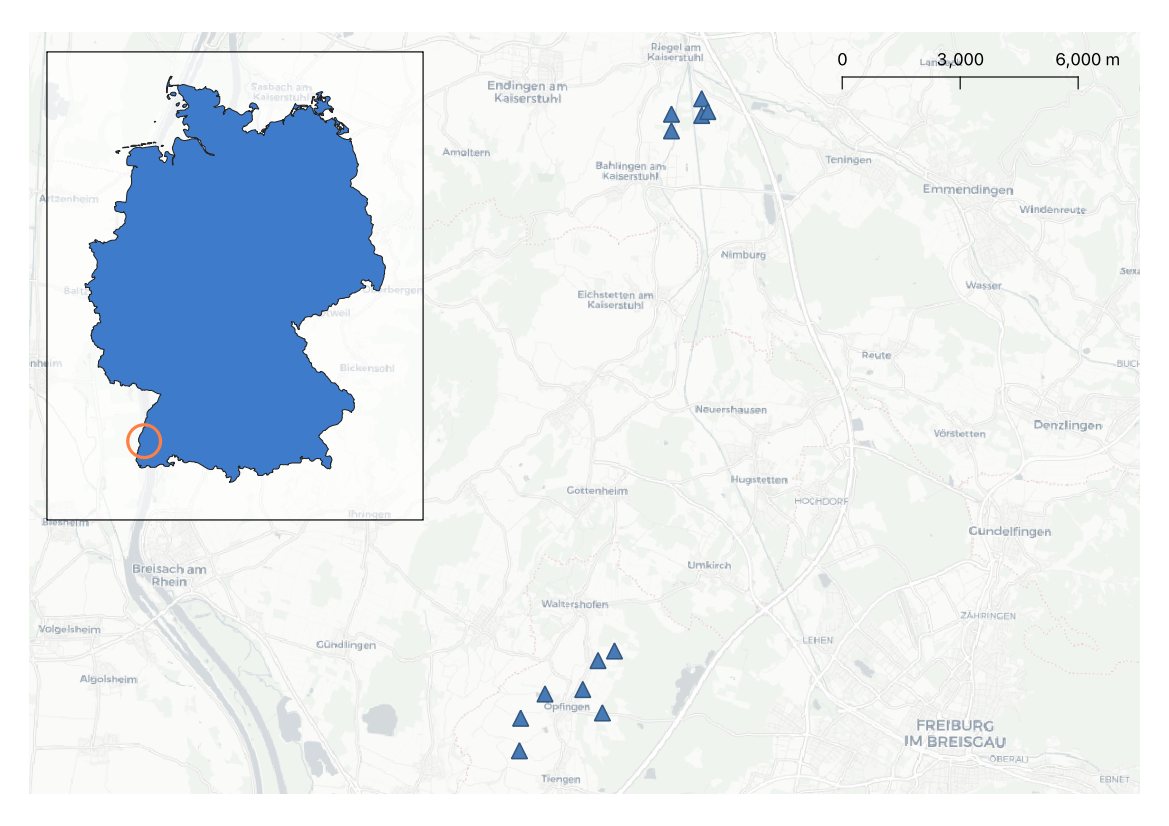

### Suppl. 2: Table of replicates

**Table S2: Number (n) of replicates per control and ploughing plot and total number of replicates, per time since establishment and sampling year.**

| **2023** | | | |
| --- | --- | --- | --- |
| time since establishment (years) | n (control plots) | n (ploughing plots) | n (total) |
| 1 | 0 | 8 | 8 |
| 2 | 4 | 0 | 4 |
| 3 | 4 | 0 | 4 |
| 4 | 2 | 2 | 4 |

| **2024** | | | |
| --- | --- | --- | --- |
| time since establishment (years) | n (control plots) | n (ploughing plots) | n (total) |
| 2 | 9 | 4 | 13 |
| 3 | 4 | 3 | 7 |
| 4 | 2 | 2 | 4 |
| 5 | 2 | 0 | 2 |

### Suppl. 3: Models

#### a) Insect biomass model

##### R Model formula: Insect biomass model

*glmmTMB(*

*daily_mass ~	(plough + sampling_year) * I(1 / time_since_establishment) +*

*(plough + sampling_year) * poly(julian_day, 2) +*

*(1 | site/trap),*

*family = Gamma(link = "log")*

*)*

##### Residual plot (DHARMa): Insect biomass model

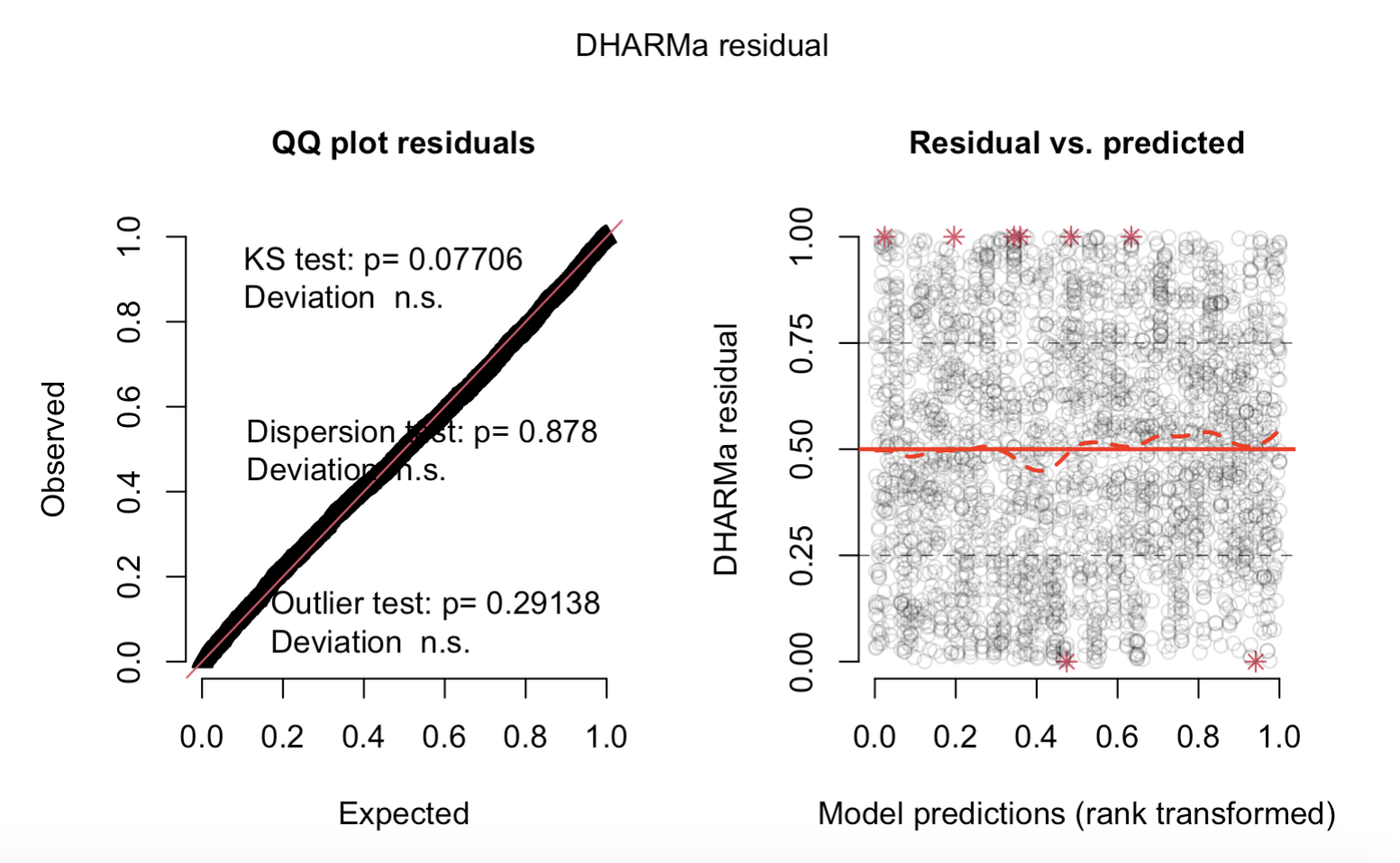

**Figure S3a: Residual plot of the R DHARMa package for the insect biomass model**

##### Coefficients: Insect biomass model

**Table S3a: Coefficient table of the insect biomass model**

| *Predictors* | *Estimates* | *std. Error* | *Statistic* | *p* |
| --- | --- | --- | --- | --- |
| (Intercept) | -2.454 | 0.267 | -9.188 | **<0.001** |
| plough [1] | -1.207 | 0.157 | -7.668 | **<0.001** |
| sampling year [2024] | -1.038 | 0.158 | -6.583 | **<0.001** |
| 1/time since  establishment | -1.864 | 0.497 | -3.752 | **<0.001** |
| julian day [1st degree] | 10.200 | 1.978 | 5.158 | **<0.001** |
| julian day [2nd degree] | -3.082 | 1.823 | -1.690 | 0.091 |
| plough [1] × 1/time since  establishment | 1.263 | 0.377 | 3.348 | **0.001** |
| sampling year [2024] ×  1/time since  establishment | 1.957 | 0.558 | 3.510 | **<0.001** |
| plough [1] × julian day  [1st degree] | -4.294 | 1.879 | -2.286 | **0.022** |
| plough [1] × julian day  [2nd degree] | 11.720 | 1.828 | 6.413 | **<0.001** |
| sampling year [2024] ×  julian day [1st degree] | 19.056 | 2.211 | 8.618 | **<0.001** |
| sampling year [2024] ×  julian day [2nd degree] | -6.623 | 2.038 | -3.250 | **0.001** |
| **Random Effects** | | | | |
| σ^2^ | 0.51 | | | |
| τ_00_ _trap:site_ | 0.09 | | | |
| τ_00_ _site_ | 0.10 | | | |
| ICC | 0.27 | | | |
| N _trap_ | 224 | | | |
| N _site_ | 12 | | | |
| Observations | 2847 | | | |
| Marginal R^2^ / Conditional R^2^ | 0.370 / 0.540 | | | |

#### b) Insect biomass model including mass occurrence samples

##### Residual plot (DHARMa): Insect biomass model including mass occurrence samples

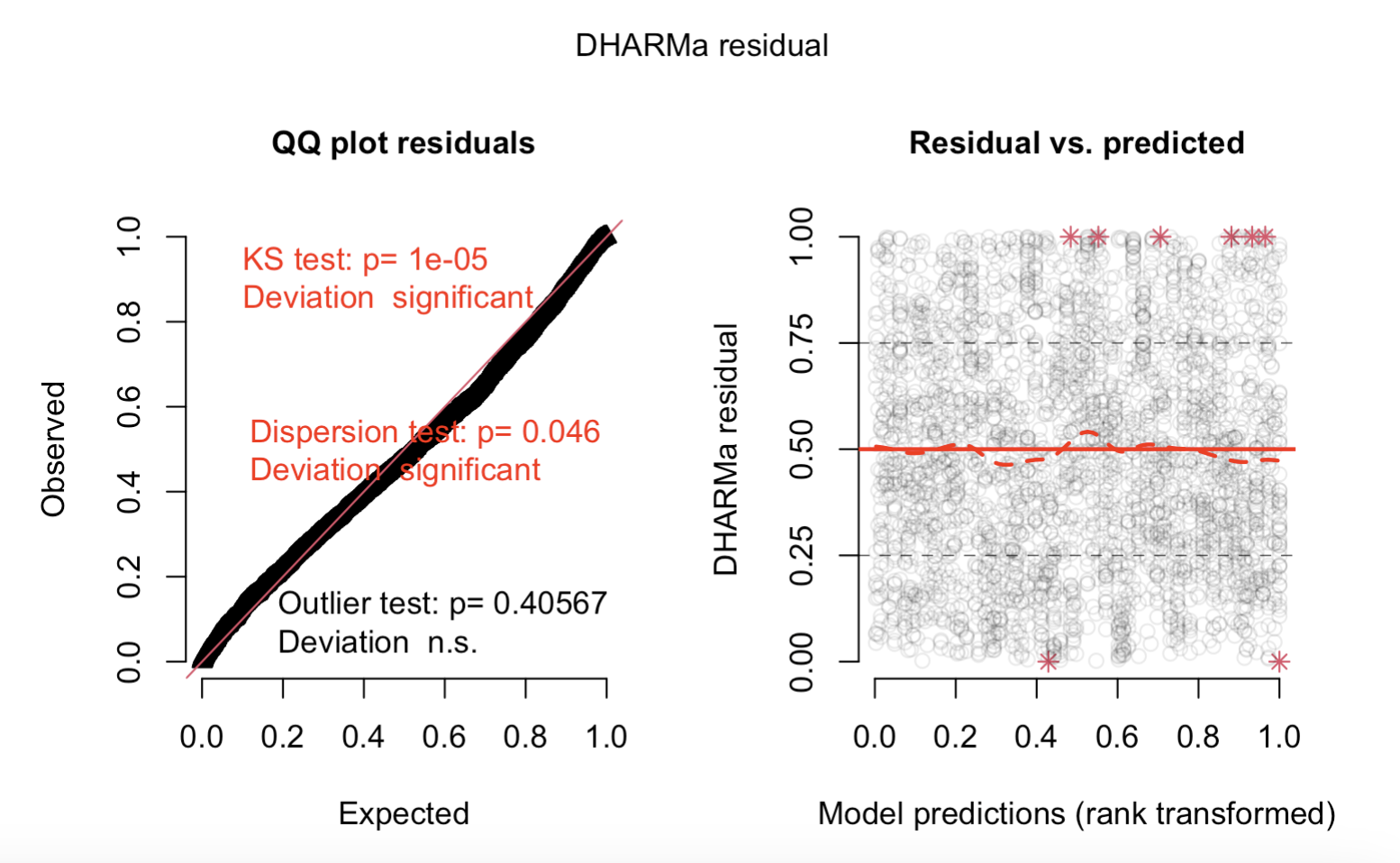

**Figure S3b: Residual plot of the R DHARMa package for the insect biomass model including mass occurrence samples**

##### Coefficients: Insect biomass model including mass occurrence samples

**Table S3b: Coefficient table of the insect biomass model including mass occurrence samples**

| *Predictors* | *Estimates* | *std. Error* | *Statistic* | *p* |
| --- | --- | --- | --- | --- |
| (Intercept) | -2.241 | 0.311 | -7.200 | **<0.001** |
| plough [1] | -1.355 | 0.177 | -7.658 | **<0.001** |
| sampling year [2024] | -1.153 | 0.175 | -6.594 | **<0.001** |
| 1/time since  establishment | -1.982 | 0.572 | -3.463 | **0.001** |
| julian day [1st degree] | 17.365 | 2.147 | 8.088 | **<0.001** |
| julian day [2nd degree] | -4.245 | 2.117 | -2.005 | **0.045** |
| plough [1] × 1/time since  establishment | 1.465 | 0.421 | 3.476 | **0.001** |
| sampling year [2024] ×  1/time since  establishment | 2.450 | 0.641 | 3.821 | **<0.001** |
| plough [1] × julian day  [1st degree] | -7.875 | 2.098 | -3.753 | **<0.001** |
| plough [1] × julian day  [2nd degree] | 11.114 | 2.057 | 5.402 | **<0.001** |
| sampling year [2024] ×  julian day [1st degree] | 22.064 | 2.413 | 9.143 | **<0.001** |
| sampling year [2024] ×  julian day [2nd degree] | -2.610 | 2.322 | -1.124 | 0.261 |
| **Random Effects** | | | | |
| σ^2^ | 0.59 | | | |
| τ_00_ _trap:site_ | 0.14 | | | |
| τ_00_ _site_ | 0.13 | | | |
| ICC | 0.32 | | | |
| N _trap_ | 224 | | | |
| N _site_ | 12 | | | |
| Observations | 2991 | | | |
| Marginal R^2^ / Conditional R^2^ | 0.391 / 0.584 | | | |

#### c) SEM Model

##### R Model formula: SEM Model

**Biomass model**

*glmmTMB(*

*sc_log_daily_mass ~ num_plough + sc_julian_day +*

*sc_log_daily_cnt_small + sc_log_daily_cnt_large +*

*(1 | trap),*

*family = gaussian()*

*)*

**Small insect model**

*glmmTMB(*

*sc_log_daily_cnt_small ~ sc_time_since_establishment +*

*sc_julian_day + num_plough +*

*(1 | site/trap),*

*family = gaussian()*

*)*

**Large insect model**

*glmmTMB(*

*sc_log_daily_cnt_large ~ sc_time_since_establishment +*

*sc_julian_day + num_plough +*

*(1 | trap),*

*family = gaussian()*

*)*

##### Coefficients: SEM model

|  | **Biomass model** | | | | **Small insects model** | | | | **Large insects model** | | | |
| --- | --- | --- | --- | --- | --- | --- | --- | --- | --- | --- | --- | --- |
| *Predictors* | *Estimates*  *[std. Est.]* | *std. Error* | *Statistic* | *p* | *Estimates*  *[std. Est.]* | *std. Error* | *Statistic* | *p* | *Estimates*  *[std. Est.]* | *std. Error* | *Statistic* | *p* |
| (Intercept) | 0.27 | 0.13 | 2.14 | **0.033** | 0.45 | 0.26 | 1.75 | 0.081 | 1.27 | 0.19 | 6.76 | **<0.001** |
| plough | -0.19  [-0.09] | 0.09 | -2.23 | **0.026** | -0.30  [-0.15] | 0.16 | -1.88 | 0.060 | -0.88  [-0.44] | 0.12 | -7.11 | **<0.001** |
| julian_day | 0.08  [0.08] | 0.03 | 3.19 | **0.001** | 0.34  [0.34] | 0.04 | 8.18 | **<0.001** | 0.35  [0.35] | 0.04 | 8.23 | **<0.001** |
| daily_cnt_small | 0.46  [0.46] | 0.03 | 14.39 | **<0.001** |  |  |  |  |  |  |  |  |
| daily_cnt_large | 0.46  [0.47] | 0.03 | 14.58 | **<0.001** |  |  |  |  |  |  |  |  |
| time_since_establishment |  |  |  |  | 0.05  [0.05] | 0.12 | 0.38 | 0.706 | 0.08  [0.08] | 0.06 | 1.24 | 0.216 |
| **Random Effects** | | | | | | | | | | | | |
| σ^2^ | 0.17 | | | | 0.56 | | | | 0.60 | | | |
| τ_00_ | 0.05 _trap_ | | | | 0.17 _trap:site_ | | | | 0.08 _trap_ | | | |
|  |  | | | | 0.13 _site_ | | | |  | | | |
| ICC | 0.21 | | | | 0.35 | | | | 0.12 | | | |
| N | 42 _trap_ | | | | 42 _trap_ | | | | 42 _trap_ | | | |
|  |  | | | | 12 _site_ | | | |  | | | |
| Observations | 326 | | | | 326 | | | | 326 | | | |
| Marginal R^2^ / Conditional R^2^ | 0.778 / 0.825 | | | | 0.141 / 0.438 | | | | 0.327 / 0.405 | | | |

**Table S3c1: Coefficient table of the SEM** **model**

*Coefficients: SEM model including mass occurrence samples*

**Table S3c2: Coefficient table of the SEM model including mass occurrence samples**

|  | **Biomass model** | | | | **Small insects model** | | | | **Large insects model** | | | |
| --- | --- | --- | --- | --- | --- | --- | --- | --- | --- | --- | --- | --- |
| *Predictors* | *Estimates*  *[std. Est.]* | *std. Error* | *Statistic* | *p* | *Estimates*  *[std. Est.]* | *std. Error* | *Statistic* | *p* | *Estimates*  *[std. Est.]* | *std. Error* | *Statistic* | *p* |
| (Intercept) | 0.23 | 0.12 | 1.86 | 0.062 | 0.48 | 0.26 | 1.89 | 0.059 | 1.19 | 0.18 | 6.48 | **<0.001** |
| plough | -0.16  [-0.08] | 0.08 | -1.96 | **0.050** | -0.33  [-0.16] | 0.16 | -2.10 | **0.036** | -0.84  [-0.41] | 0.12 | -6.85 | **<0.001** |
| julian_day | 0.09  [0.09] | 0.02 | 3.43 | **0.001** | 0.34  [0.34] | 0.04 | 8.54 | **<0.001** | 0.40  [0.40] | 0.04 | 9.68 | **<0.001** |
| daily_cnt_small | 0.42  [0.42] | 0.03 | 14.28 | **<0.001** |  |  |  |  |  |  |  |  |
| daily_cnt_large | 0.51  [0.51] | 0.03 | 16.88 | **<0.001** |  |  |  |  |  |  |  |  |
| time_since_establishment |  |  |  |  | 0.03  [0.78] | 0.12 | 0.27 | 0.784 | 0.08  [0.08] | 0.06 | 1.40 | 0.162 |
| **Random Effects** | | | | | | | | | | | | |
| σ^2^ | 0.15 | | | | 0.54 | | | | 0.57 | | | |
| τ_00_ | 0.04 _trap_ | | | | 0.17 _trap:site_ | | | | 0.08 _trap_ | | | |
|  |  | | | | 0.13 _site_ | | | |  | | | |
| ICC | 0.23 | | | | 0.35 | | | | 0.12 | | | |
| N | 42 _trap_ | | | | 42 _trap_ | | | | 42 _trap_ | | | |
|  |  | | | | 12 _site_ | | | |  | | | |
| Observations | 336 | | | | 336 | | | | 336 | | | |
| Marginal R^2^ / Conditional R^2^ | 0.798 / 0.843 | | | | 0.150 / 0.450 | | | | 0.350 / 0.429 | | | |
